# Synergy and remarkable specificity of antimicrobial peptides *in vivo* using a systematic knockout approach

**DOI:** 10.1101/493817

**Authors:** M.A. Hanson, A. Dostálová, C. Ceroni, M. Poidevin, S. Kondo, B. Lemaitre

## Abstract

Antimicrobial peptides (AMPs) are host-encoded antibiotics that combat invading microorganisms. These short, cationic peptides have been implicated in many biological processes, primarily involving innate immunity. *In vitro* studies have shown AMPs kill bacteria and fungi at physiological concentrations, but little validation has been done *in vivo.* We utilised CRISPR gene editing to delete all known immune inducible AMPs of *Drosophila,* namely: 4 Attacins, 4 Cecropins, 2 Diptericins, Drosocin, Drosomycin, Metchnikowin and Defensin. Using individual and multiple knockouts, including flies lacking all 14 AMP genes, we characterize the *in vivo* function of individual and groups of AMPs against diverse bacterial and fungal pathogens. We found that *Drosophila* AMPs act primarily against Gram-negative bacteria and fungi, acting either additively or synergistically. We also describe remarkable specificity wherein certain AMPs contribute the bulk of microbicidal activity against specific pathogens, providing functional demonstrations of highly specific AMP-pathogen interactions in an *in vivo* setting.

## Introduction

While innate immune mechanisms were neglected during the decades where adaptive immunity captured most of the attention, they have become central to our understanding of immunology. Recent emphasis on innate immunity has, however, mostly focused on the first two phases of the immune response: microbial recognition and associated downstream signaling pathways. In contrast, how innate immune effectors individually or collectively contribute to host resistance has not been investigated to the same extent. The existence of multiple effectors that redundantly contribute to host resistance has hampered their functional characterization by genetic approaches^1^. The single mutation methodology that still prevails today has obvious limits in the study of immune effectors, which often belong to large gene families. As such, our current understanding of the logic underlying the roles of immune effectors is only poorly defined. As a consequence, the key parameters that influence host survival associated with a successful immune response are not well characterized. In this paper, we harnessed the power of the CRISPR gene editing approach to study the function of *Drosophila* antimicrobial peptides in host defence both individually and collectively.

Antimicrobial peptides (AMPs) are small, cationic, usually amphipathic peptides that contribute to innate immune defence in plants and animals^2–4^. They display potent antimicrobial activity *in vitro* by disrupting negatively-charged microbial membranes, but AMPs can also target specific microbial processes^5–7^. Their expression is induced to very high levels upon challenge to provide microbicidal concentrations in the μM range. Numerous studies have revealed unique roles that AMPs may play in host physiology, including anti-tumour activity^8,9^, inflammation in aging^10–12^, involvement in memory^13,14^, mammalian immune signaling^15,16^, wound-healing^17,18^, regulation of the host microbiota^19,20^, tolerance to oxidative stress^21,22^, and of course microbicidal activity^1,2,23^. The fact that AMP genes are immune inducible and expressed at high levels has led to the common assumption they play a vital role in the innate immune response^24^. However, little is known in most cases about how AMPs individually or collectively contribute to animal host defence. *In vivo* functional analysis of AMPs has been hampered by the sheer number and small size of these genes, making them difficult to mutate with traditional genetic tools (but *e.g.* see^25,26^).

Since the first animal AMPs were discovered in silk moths^27^, insects and particularly *Drosophila melanogaster* have emerged as a powerful model for characterizing their function. There are currently seven known families of inducible AMPs in *D. melanogaster.* Their activities have been determined either *in vitro* by using peptides directly purified from flies or produced in heterologous systems, or deduced by comparison with homologous peptides isolated in other insect species: Drosomycin and Metchnikowin show antifungal activity^28,29^; Cecropins (four inducible genes) and Defensin have both antibacterial and some antifungal activities^30–33^; and Drosocin, Attacins (four genes) and Diptericins (two genes) primarily exhibit antibacterial activity^6,34–37^. In *Drosophila,* these AMPs are produced either locally at various surface epithelia in contact with environmental microbes^38–40^, or secreted systemically into the hemolymph, the insect blood. During systemic infection, these 14 antimicrobial peptides are strongly induced in the fat body, an organ analogous to the mammalian liver.

The systemic production of AMPs is regulated at the transcriptional level by two NF-kB pathways, the Toll and Imd pathways, which are activated by different classes of microbes. The Toll pathway is predominantly responsive to Gram-positive bacteria and fungi, and accordingly plays a major role in defence against these microbes. In contrast, the Imd pathway is activated by Gram-negative bacteria and a subset of Gram-positive bacteria with DAP-type peptidoglycan, and mutations affecting this pathway cause profound susceptibility to Gram-negative bacteria^41,42^. However, the expression pattern of AMP genes is complex as each gene is expressed with different kinetics and can often receive transcriptional input from both pathways^42,43^. This ranges from *Diptericin,* which is tightly regulated by the Imd pathway, to *Drosomycin,* whose expression is mostly regulated by the Toll pathway^41^, except at surface epithelia where *Drosomycin* is under the control of Imd signaling^44^. While a critical role of AMPs in *Drosophila* host defence is supported by transgenic flies overexpressing a single AMP^33^, the specific contributions of each of these AMPs has not been tested. Indeed loss-of-function mutants for most AMP genes were not previously available due to their small size, making them difficult to mutate before the advent of CRISPR/Cas9 technology. Despite this, the great susceptibility to infection of mutants with defective Toll and Imd pathways is commonly attributed to the loss of the AMPs they regulate, though these pathways control hundreds of genes awaiting characterization^42^. Strikingly, Clemmons *et* al.^45^ recently reported that flies lacking a set of uncharacterized Toll-responsive peptides (named Bomanins) succumb to infection by Gram-positive bacteria and fungi at rates similar to Toll-deficient mutants^45^. This provocatively suggests that Bomanins, and not AMPs, might be the predominant effectors downstream of the Toll pathway; yet synthesized Bomanins do not display antimicrobial activity *in vitro^46^.* Thus, while today the fly represents one of the best-characterized animal immune systems, the contribution of AMPs as immune effectors is poorly defined as we still do not understand why Toll and Imd pathway mutants succumb to infection.

In this paper, we took advantage of recent gene editing technologies to delete each of the known immune inducible AMP genes of *Drosophila.* Using single and multiple knockouts, as well as a variety of bacterial and fungal pathogens, we have characterized the *in vivo* function of individual and groups of antimicrobial peptides. We reveal that AMPs can play highly specific roles in defence, being vital for surviving certain infections yet dispensable against others. We highlight key interactions amongst immune effectors and pathogens and reveal to what extent these defence peptides act in concert or alone.

## Results

### Generation and characterization of AMP mutants

We generated null mutants for the fourteen *Drosophila* antimicrobial peptide genes that are induced upon systemic infection. These include five single gene mutations affecting *Defensin (Def^SK3^), Attacin C* (*AttC^Mi^*), *Metchnikowin (Mtk^R1^), Attacin D* (*AttD^SK1^*) and *Drosomycin (Drs^R1^)* respectively, and three small deletions removing both *Diptericins DptA* and *DptB (Dpt^SK1^),* the four *Cecropins CecA1, CecA2, CecB, and CecC* (*Cec^SK6^*) and the gene cluster containing *Drosocin,* and *Attacins AttA* & *AttB (Dro-AttAB^SK2^).* All mutations/deletions were made using the CRISPR editing approach with the exception of *Attacin C,* which was disrupted by insertion of a *Minos* transposable element^47^, and the *Drosomycin* and *Metchnikowin* deletions generated by homologous recombination (Fig. 1A). To disentangle the role of *Drosocin* and *AttA/AttB* in the *Dro-AttAB^SK2^* deletion, we also generated an individual *Drosocin* mutant (Dro^SK4^); for complete information, see Figure S1. We then isogenized these mutations for at least seven generations into the *w^1118^* DrosDel isogenic genetic background^48^ (iso w^1118^). Then, we recombined these eight independent mutations into a background lacking all 14 inducible AMPs referred to as *“ΔAMPs.” ΔAMPs* flies were viable and showed no morphological defects. To confirm the absence of AMPs in our *ΔAMPs* background, we performed a MALDI-TOF analysis of hemolymph from both unchallenged and immune-challenged flies infected by a mixture of *Escherichia coli* and *Micrococcus luteus.* This analysis revealed the presence of peaks induced upon challenge corresponding to AMPs in wild-type but not *ΔAMPs* flies. Importantly it also confirmed that induction of most other immune-induced molecules (IMs)^49^, was unaffected in *ΔAMPs* flies (Fig. 1B). Of note, we failed to observe two IMs, IM7 and IM21, in our *ΔAMPs* flies, suggesting that these unknown peptides are secondary products of AMP genes. We further confirmed that Toll and Imd NF-ĸB signaling pathways were intact in *ΔAMPs* flies by measuring the expression of target genes of these pathways (Fig. 1C-D). This demonstrates that *Drosophila* AMPs are not signaling molecules required for Toll or Imd pathway activity. We also assessed the role of AMPs in the melanization response, wound clotting, and hemocyte populations. After clean injury, *ΔAMPs* flies survive as wild-type (Fig. 1 supplement A). We found no defect in melanization (χ^2^, p = .34, Fig. 1 supplement B) as both adults and larvae strongly melanize the cuticle following clean injury, (Fig. 1 supplement C). Furthermore, we visualized the formation of clot fibers *ex vivo* using the hanging drop assay and PNA staining^50^ in hemolymph of both wild-type and *ΔAMPs* larvae (Fig. 1 supplement D). Hemocyte counting (i.e. crystal cells, FACS) did not reveal any deficiency in hemocyte populations of *ΔAMPs* larvae (Fig. 1 supplement E, F, and not shown). Altogether, our study suggests that *Drosophila* AMPs are primarily immune effectors, and not regulators of innate immunity.

**Figure 1:**
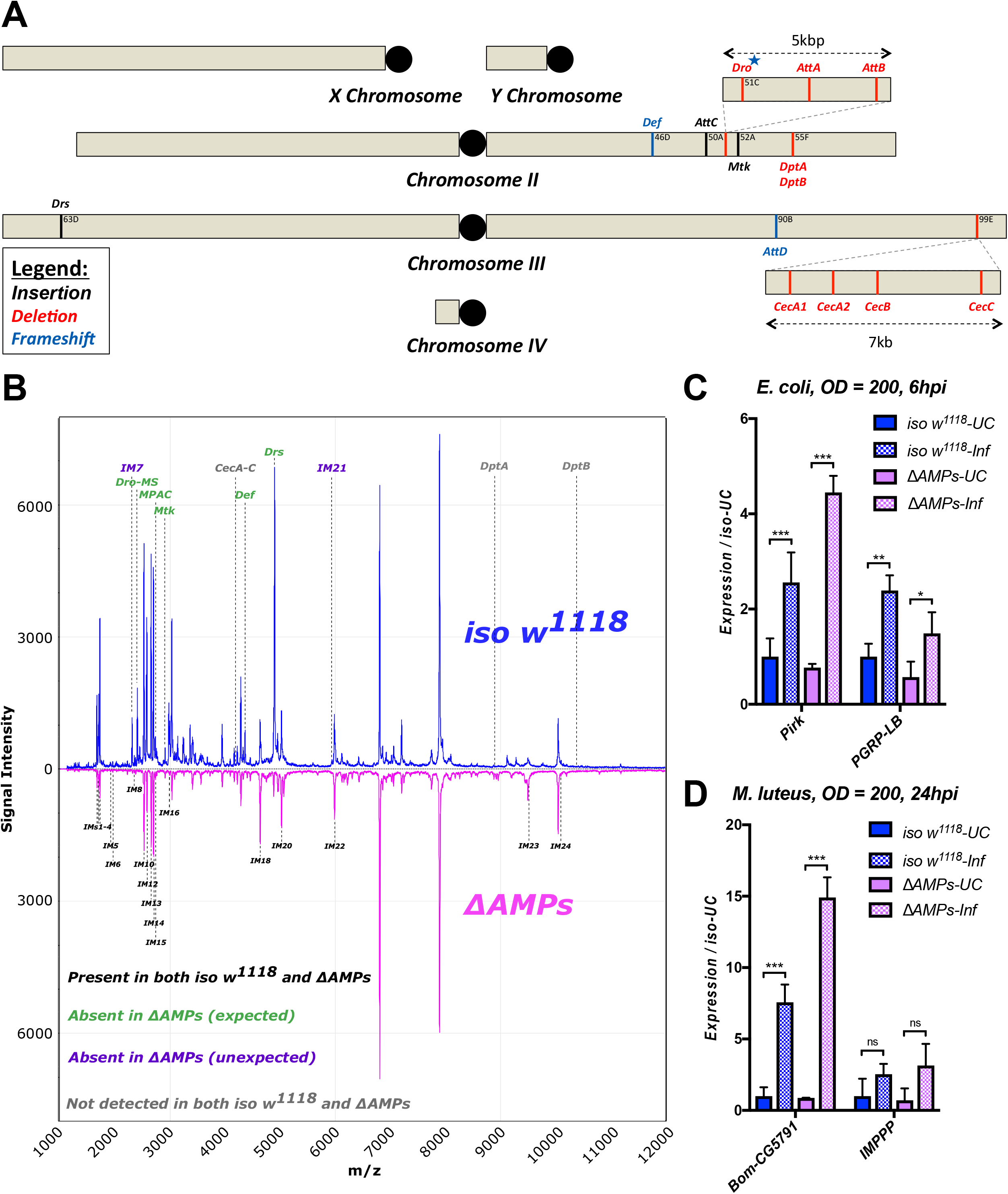
Description of *AMP* mutants. A) Chromosomal location of *AMP* genes that were deleted. Each mutation is color-coded with the mutagenic agent: black, a *Minos* insertion or homologous recombination, red, CRISPR-CAS9 mediated deletion, and blue CRISPR CAS9 mediated indel causing a nonsense peptide. B) A representative MALDI-TOF analysis of hemolymph samples from immune-challenged (1:1 *E. coli* and *M. luteus* at 0D600 = 200) *iso w^1118^* and *ΔAMPs* flies as described in Üttenweiller-Joseph et al.^49^. No AMP-derived products were detected in the hemolymph samples of *ΔAMPs* flies. No signals for IM7, nor IM21 were observed in the hemolymph samples of *ΔAMPs* mutants suggesting that these uncharacterized immune-induced molecules are the products of AMP genes. The Imd pathway (C) and Toll pathway (D) are active and respond to immune challenge in *ΔAMPs* flies. We used alternate readouts to monitor the Toll and Imd pathways: *pirk* and *PGRP-LB* for Imd pathway and *CG5791 (Bomanin)* and *IMPPP* for Toll signaling^42,72^. UC = unchallenged, Inf = infected. hpi = hours post-infection. Expression normalized with *iso w^1118^-UC* set to a value of 1.

### AMPs are essential for combating Gram-negative bacterial infection

We used these *ΔAMPs* flies to explore the role that AMPs play in defence against pathogens during systemic infection. We first focused our attention on Gram-negative bacterial infections, which are combatted by Imd pathway-mediated defence in *Drosophila*^1^. We challenged wild-type and *ΔAMPs* flies with six different Gram-negative bacterial species, using inoculation doses (given as 0D600) selected such that at least some wild-type flies were killed (Fig. 2). In our survival experiments, we also include *Relish* mutants *(Rel^E20^)* that lack a functional Imd response and are known to be very susceptible to this class of bacteria^51^. Globally, *ΔAMPs* flies were extremely susceptible to all Gram-negative pathogens tested (Fig. 2, light blue plots). The susceptibility of AMP-deficient flies to Gram-negative bacteria largely mirrored that of *Rel^E20^* flies. For all Gram-negative infections tested, *ΔAMPs* flies show a higher bacterial count at 18 hours post-infection (hpi) indicating that AMPs actively inhibit bacterial growth, as expected of ‘antimicrobial peptides’ (Fig. 2 supplement A). Use of GFP-expressing bacteria show that bacterial growth in *ΔAMPs* flies radiates from the wound site until it spreads systemically (Fig. 2 supplement B,C). Collectively, the use of AMP-deficient flies reveals that AMPs are major players in resistance to Gram-negative bacteria, and likely constitute an essential component of the Imd pathway’s contribution for survival against these germs.

**Figure 2:**
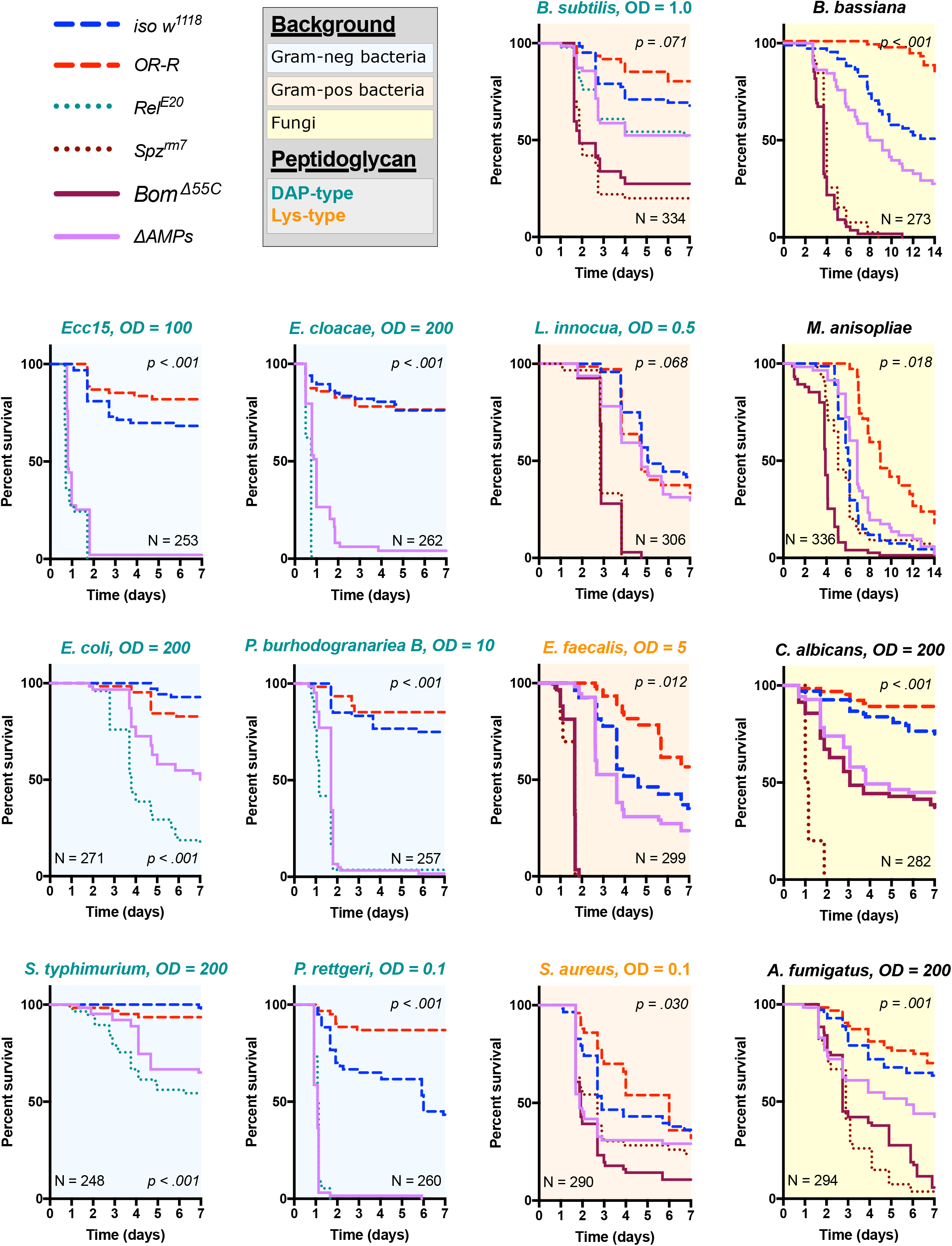
Survival of *ΔAMPs* flies to diverse microbial challenges. Control lines for survival experiments included two wild-types *(w;Drosdel (iso w^1118^)* and Oregon R *(OR-R)* as an alternate wild-type), mutants for the Imd response *(Rel^E20^),* mutants for Toll signaling (spz^rm7^), and mutants for Bomanins *(Bom^Δ55C^). ΔAMPs* flies are extremely susceptible to infection with Gram-negative bacteria (blue backgrounds). Unexpectedly, *ΔAMPs* flies were not markedly susceptible to infection with Gram-positive bacteria (orange backgrounds), while *Bom^Δ55C^* flies were extremely susceptible, often mirroring *spz^rm7^* mutants. This pattern of *Bom^Δ55C^* susceptibility held true for fungal infections (yellow backgrounds). *ΔAMPs* flies are somewhat susceptible to fungal infections, but the severity shifts with different fungi. Pellet densities are reported for all systemic infections in OD at 600nm. P-values are given for *ΔAMPs* flies compared to *iso w^1118^* using a Cox-proportional hazards model.

### Bomanins and to a lesser extent AMPs contribute to resistance against Gram-positive bacteria and fungi

Previous studies have shown that resistance to Gram-positive bacteria and fungi in *Drosophila* is mostly mediated by the Toll pathway, although the Imd pathway also contributes to some extent^41,43,52,53^. Moreover, a deletion removing eight uncharacterized Bomanins *(Bom^Δ55C^)* induces a strong susceptibility to both Gram-positive bacteria and fungi^45^, suggesting that Bomanins are major players downstream of Toll in the defence against these germs. This prompted us to explore the role of antimicrobial peptides in defence against Gram-positive bacteria and fungi. We first challenged wild-type and *ΔAMPs* flies with two lysine-type (E. *faecalis, S. aureus)* and two DAP-type peptidoglycan containing Gram-positive bacterial species *(B. subtilis, L. innocua).* We observed that *ΔAMPs* flies display only weak or no increased susceptibility to infection with these Gram-positive bacterial species, as *ΔAMPs* survival rates were closer to the wild-type than to *späztle* flies *(spz^rm7^)* lacking a functional Toll pathway (Fig. 2, orange plots). Meanwhile, *Bom^Δ55C^* mutants consistently phenocopied *spz^rm7^* flies, confirming the important contribution of these peptides in defence against Gram-positive bacteria^45^.

Next, we monitored the survival of *ΔAMPs* to the yeast *Candida albicans,* the opportunistic fungus *Aspergillus fumigatus* and two entomopathogenic fungi, *Beauveria bassiana,* and *Metarhizium anisopliae.* For the latter two, we used a natural mode of infection by spreading spores on the cuticle^41^. *ΔAMPs* flies were more susceptible to fungal infections with *B. bassiana, A. fumigatus,* and *C. albicans,* but not *M. anisopliae* (Fig. 2, yellow plots). In all instances, *Bom^Δ55C^* mutants were as or more susceptible to fungal infection than *ΔAMPs* flies, approaching Toll-deficient mutant levels. Collectively, our data demonstrate that AMPs are major immune effectors in defence against Gram-negative bacteria and have a less essential role in defence against bacteria and fungi.

### A combinatory approach to explore AMP interactions

The impact of the *ΔAMPs* deletion on survival could be due to the action of certain AMPs having a specific effect, or more likely due to the combinatory action of coexpressed AMPs. Indeed, cooperation of AMPs to potentiate their microbicidal activity has been suggested by numerous *in vitro* approaches^7,54,55^, but rarely in an *in vivo* context^56^. Having shown that AMPs as a whole significantly contribute to fly defence, we next explored the contribution of individual peptides to this effect. To tackle this question in a systematic manner, we performed survival analyses using fly lines lacking one or several AMPs, focusing on pathogens with a range of virulence that we previously showed to be sensitive to the action of AMPs. This includes the yeast *C. albicans* and the Gram-negative bacterial species *P. burhodogranariea, P. rettgeri, Ecc15,* and *E. cloacae.* Given eight independent AMP mutations, over 250 combinations of mutants are possible, making a systematic analysis of AMP interactions a logistical nightmare. Therefore, we designed an approach that would allow us to characterize their contributions to defence by deleting groups of AMPs. To this end, we generated three groups of combined mutants: flies lacking the primarily antibacterial *Defensin* and *Cecropins* (Group A, mostly regulated by the Imd pathway), flies lacking the antibacterial Proline-rich *Drosocin,* and the antibacterial Glycine-rich *Diptericins* and *Attacins* (Group B, regulated by the Imd pathway), and flies lacking the two antifungal peptide genes *Metchnikowin* and *Drosomycin* (Group C, mostly regulated by the Toll pathway). We then combined these three groups to generate flies lacking AMPs from groups A and B (AB), A and C (AC), or B and C (BC). Finally, flies lacking all three groups are our *ΔAMPs* flies, which are highly susceptible to a number of infections. By screening these seven genotypes as well as individual mutants, we were able to assess potential interactions between AMPs of different groups, as well as decipher the function of individual AMPs.

### Drosomycin and Metchnikowin additively contribute to defence against the yeast C. albicans

We first applied this AMP-groups approach to infections with the relatively avirulent yeast *C. albicans.* Previous studies have shown that Toll, but not Imd, contributes to defence against this fungus^57,58^. Thus, we suspected that the two antifungal peptides, Drosomycin and Metchnikowin, could play a significant role in the susceptibility of *ΔAMPs* flies to this yeast. Consistent with this, Group C flies lacking *Metchnikowin* and *Drosomycin* were more susceptible to infection (p < .001 relative to *iso w^1118^)* with a survival rate similar to *ΔAMPs* flies (Fig. 3A). Curiously, AC deficient flies that also lack *Cecropins* and *Defensin* survived better than Group C deficient flies (Log-Rank p = .014). We have no explanation for this interaction, but this could be due to i) a better canalization of the immune response by preventing the induction of ineffective AMPs, ii) complex biochemical interactions amongst the AMPs involved, or iii) differences in genetic background generated by additional recombination. We then investigated the individual contributions of *Metchnikowin* and *Drosomycin* to survival to *C. albicans.* We found that both *Mtk^R1^* and *Drs^R1^* individual mutants were somewhat susceptible to infection, but notably only *Mtk; Drs* compound mutants reached *ΔAMPs* levels of susceptibility (Fig. 3B). This cooccurring loss of resistance appears to be primarily additive (Mutant, Cox Hazard Ratio (HR), p-value: *Mtk^R1^, HR = +1.17, p = .008; Drs^R1^, HR = +1.85, p < .001; Mtk*Drs, HR = -0.80, p =.116).* We observed that Group C deficient flies eventually succumb to uncontrolled *C. albicans* growth by monitoring yeast titre, indicating that these AMPs indeed act by suppressing yeast growth (Fig. 3C).

**Figure 3:**
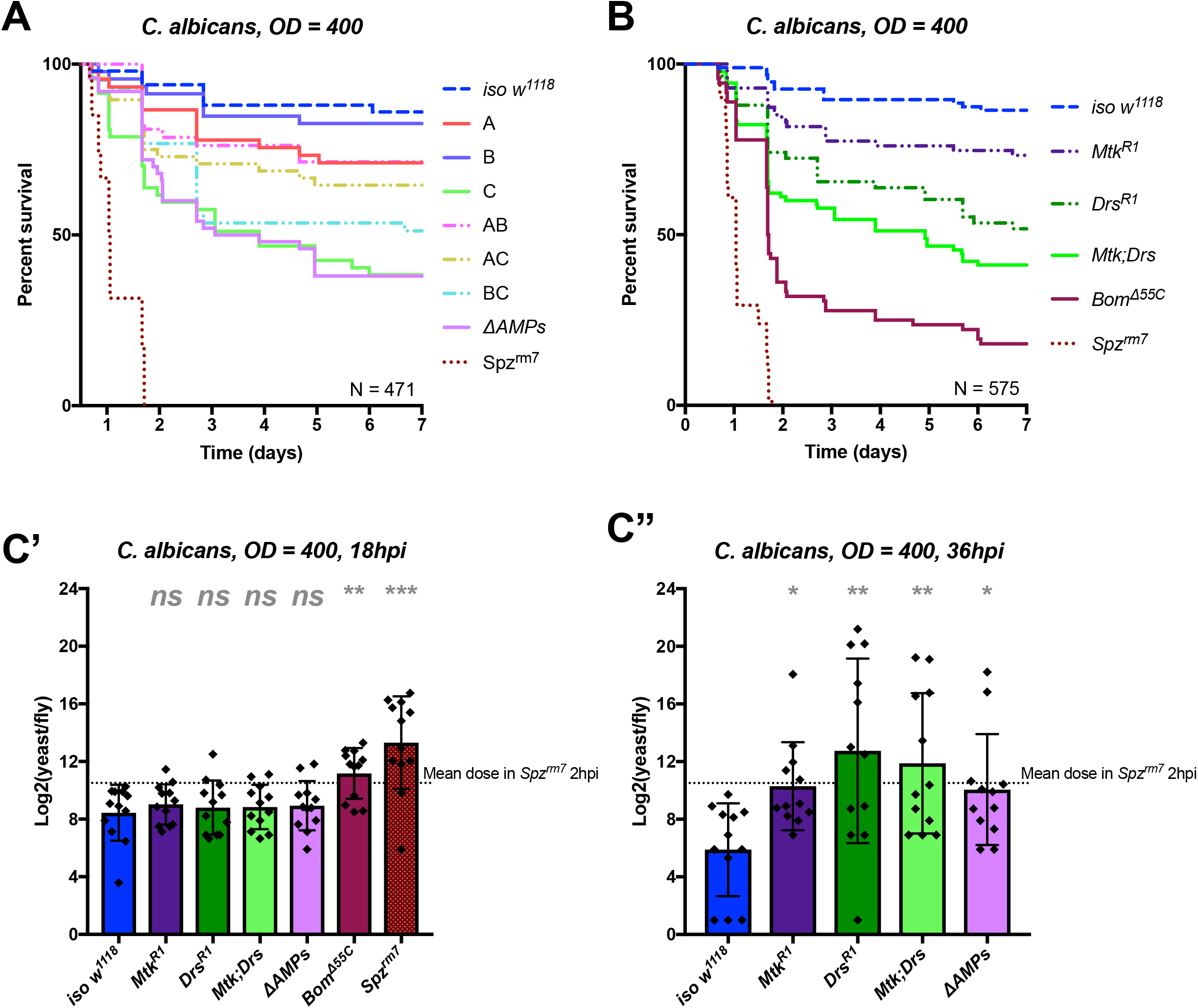
Identification of AMPs involved in the susceptibility of *ΔAMPs* flies to *C. albicans.* A) Survival of mutants for groups of AMPs reveals that loss of only Toll-responsive Group C peptides (Metchnikowin and Drosomycin) is required to recapitulate the susceptibility of *ΔAMPs* flies. Co-occurring loss of groups A and C has a net protective effect (A*C: HR = -1.71, p = .002). B) Further dissection of Group C mutations reveals that both Metchnikowin and Drosomycin contribute to resist *C. albicans* survival (p = .008 and p < .001 respectively). The interaction of Metchnikowin and Drosomycin was not different from the sum of their individual effects *(Mtk*Drs:* HR = -0.80, p = .116). Fungal loads of individual flies at 18hpi. At this time point, *Bom^Δ55C^* mutants and *spz^rm7^* flies have already failed to constrain *C. albicans* growth (C’). Fungal titres at 36hpi (C”), a time point closer to mortality for many AMP mutants, show that some AMP mutants fail to control fungal load, while wild-type flies consistently controlled fungal titre. One-way ANOVA: not significant = *ns,* p < .05 = *, p < .01 = **, and p < .001 = *** relative to *iso w^1118^*.

In conclusion, our study provides an *in vivo* validation of the potent antifungal activities of Metchnikowin and Drosomycin^28,29^, and highlights a clear example of additive cooperation of AMPs.

### AMPs synergistically contribute to defence against P. burhodogranariea

We next analyzed the contribution of AMPs in resistance to infection with the moderately virulent Gram-negative bacterium *P. burhodogranariea.* We found that Group B mutants lacking *Drosocin,* the two *Diptericins,* and the four *Attacins,* were as susceptible to infection as *ΔAMPs* flies (Fig. 4A), while flies lacking the antifungal peptides Drosomycin and Metchnikowin (Toll-regulated, Group C) resisted the infection as wild-type. Flies lacking *Defensin* and the four *Cecropins* (Group A) showed an intermediate susceptibility, but behave as wild-type in the additional absence of Toll Group C peptides (Group AC). Thus, we again observed a better survival rate with the co-occurring loss of Group A and C peptides (see possible explanation above). In this case Group A flies were susceptible while AC flies were not. Flies individually lacking *Defensin* or the four *Cecropins* were weakly susceptible to *P. burhodogranariea* (p = .022 and p = 0.040 respectively), however the interaction term between *Defensin and* the *Cecropins* was not significant *(Def^SK3^*Cec^SK6^*, HR = -0.28, p = .382), indicating the susceptibility of Group A flies arises from additive loss of resistance (Figure 4 supplement A).

**Figure 4:**
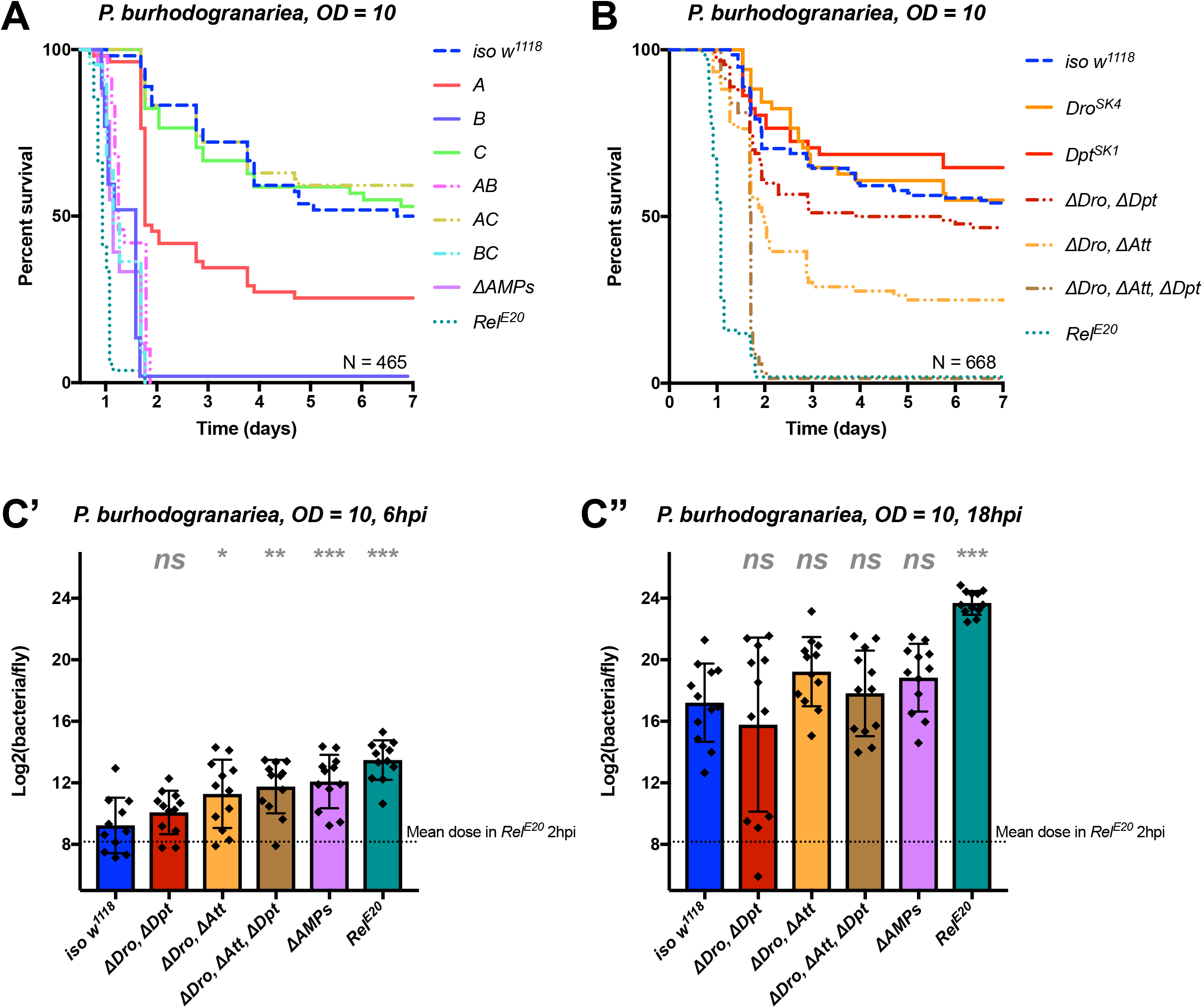
Identification of AMPs involved in the susceptibility of *ΔAMPs* flies to *P. burhodogranariea.* A) Survival of mutants for groups of AMPs reveals that loss of Imd-responsive Group B peptides (Drosocin, Attacins, and Diptericins) recapitulates the susceptibility of *ΔAMPs* flies. Loss of Group A peptides also resulted in strong susceptibility (p < .001) due to additive effects of Defensin and Cecropins (Fig. 4 supplement). B) Further dissection of AMPs deleted in Group B reveals that only the loss of all Drosocin, Attacin, and Diptericin gene families leads to susceptibility similar to *ΔAMPs* flies. Simultaneous loss of *Attacins* and *Diptericins* results in a synergistic loss of resistance *(ΔAtt*ΔDpt:* HR = +1.45, p < .001). C) Bacterial loads of individual flies at 6hpi (C’). At this time point, most AMP mutants had significantly higher bacterial loads compared to wild-type flies. At 18hpi (C”), differences in bacterial load are reduced, likely owing to the high chronic load *P. burhodogranariea* establishes even in surviving flies^24^. Meanwhile *Rel^E20^* flies succumb ~18 hours earlier than *ΔAMPs* flies in survival experiments, and already have significantly higher loads. One-way ANOVA: not significant = *ns*, p < .05 = *, p < .01 = **, and p < .001 = *** relative to *iso w*^1118^.

Following the observation that Group B flies were as susceptible as *ΔAMPs* flies, we sought to better decipher the contribution of each Group B AMP to resistance to *P. burhodogranariea.* We observed that mutants for *Drosocin* alone *(Dro^SK4^),* or the *DiptericinA/B* deficiency were not susceptible to this bacterium (Fig. 4B). We additionally saw no marked susceptibility of *Drosocin-Attacin A/B* deficient flies, nor *Attacin C* or *Attacin D* mutants (not shown). Interestingly, we found that compound mutants lacking *Drosocin* and *Attacins A, B, C,* and *D* (Fig. 4B: *‘ΔDro, ΔAtt’*), or *Drosocin* and *Diptericins DptA* and *DptB (‘ΔDro, ΔDp’)* displayed an intermediate susceptibility. Only the Group B mutants lacking *Drosocin,* all *Attacins,* and both *Diptericins (ΔDro, ΔAtt, ΔDpt)* phenocopied *ΔAMPs* flies (Fig. 4B), with synergistic interactions observed upon co-occurring loss of *Attacins* and *Diptericins (ΔAtt*ΔDpt:* HR = +1.45, p < .001). By 6hpi, bacterial titres of individual flies already showed significant differences in the most susceptible genotypes (Fig. 4C), though these differences were reduced by 18hpi likely owing to the high chronic load *P. burhodogranariea* establishes in surviving flies^24^; also see Fig. 2 supplement A.

Collectively, the use of various compound mutants reveals that several Imd-responsive AMPs, notably Drosocin, Attacins, and Diptericins, jointly contribute to defence against *P. burhodogranariea* infection. A strong susceptibility of Group B flies was also observed upon infection with *Ecc15,* another Gram-negative bacterium commonly used to infect flies^59^ (Fig. 4 supplement B).

### *Diptericins* alone contribute to defence against *P. rettgeri*

We continued our exploration of AMP interactions using our AMP groups approach with the fairly virulent *P. rettgeri* (strain Dmel), a strain isolated from wild-caught *Drosophila* hemolymph^60^. We were especially interested by this bacterium as previous studies^61,62^ have shown a correlation between susceptibility to *P. rettgeri* and a polymorphism in the *Diptericin A* gene pointing to a specific AMP-pathogen interaction. Use of compound mutants revealed only loss of Group B AMPs was needed to reach the susceptibility of *ΔAMPs* and *Rel^E20^* flies (Fig. 5A). Use of individual mutant lines however revealed a pattern strikingly different from that *P. burhodogranariea,* as the sole *Diptericin A/B* deficiency caused susceptibility similar to Group B, *ΔAMPs,* and *Rel^E20^* flies (Fig. 5B,C). We further confirmed this susceptibility using a *DptA RNAi* construct (Fig. 5 supplement A, B). Moreover, flies carrying the *Dpt^SK1^* mutation over a deficiency *(Df(2R)Exel6067)* were also highly susceptible to *P. rettgeri* (Fig. 5D). Interestingly, flies that were heterozygotes for *Dpt^SK1^* or the *Df(2R)Exel6067* that have only one copy of the two *Diptericins* were markedly susceptible to infection with *P. rettgeri* (Fig. 5D). This indicates that a full transcriptional output of *Diptericin* is required over the course of the infection to resist *P. rettgeri* infection (Fig. 5E). Altogether, our results suggest that only the *Diptericin* gene family, amongst the many AMPs regulated by the Imd pathway, provides the full AMP-based contribution to defence against this bacterium. To test this hypothesis, we generated a fly line lacking all the AMPs except *DptA* and *DptB (ΔAMPs+^Dpt^).* Strikingly, *ΔAMPs+^Dpt^* flies have the same survival rate as wild-type flies, further emphasizing the specificity of this interaction (Fig. 5B). Bacterial counts confirm that the susceptibility of these *Diptericin* mutants arises from an inability of the host to suppress bacterial growth (Fig. 5C).

**Figure 5:**
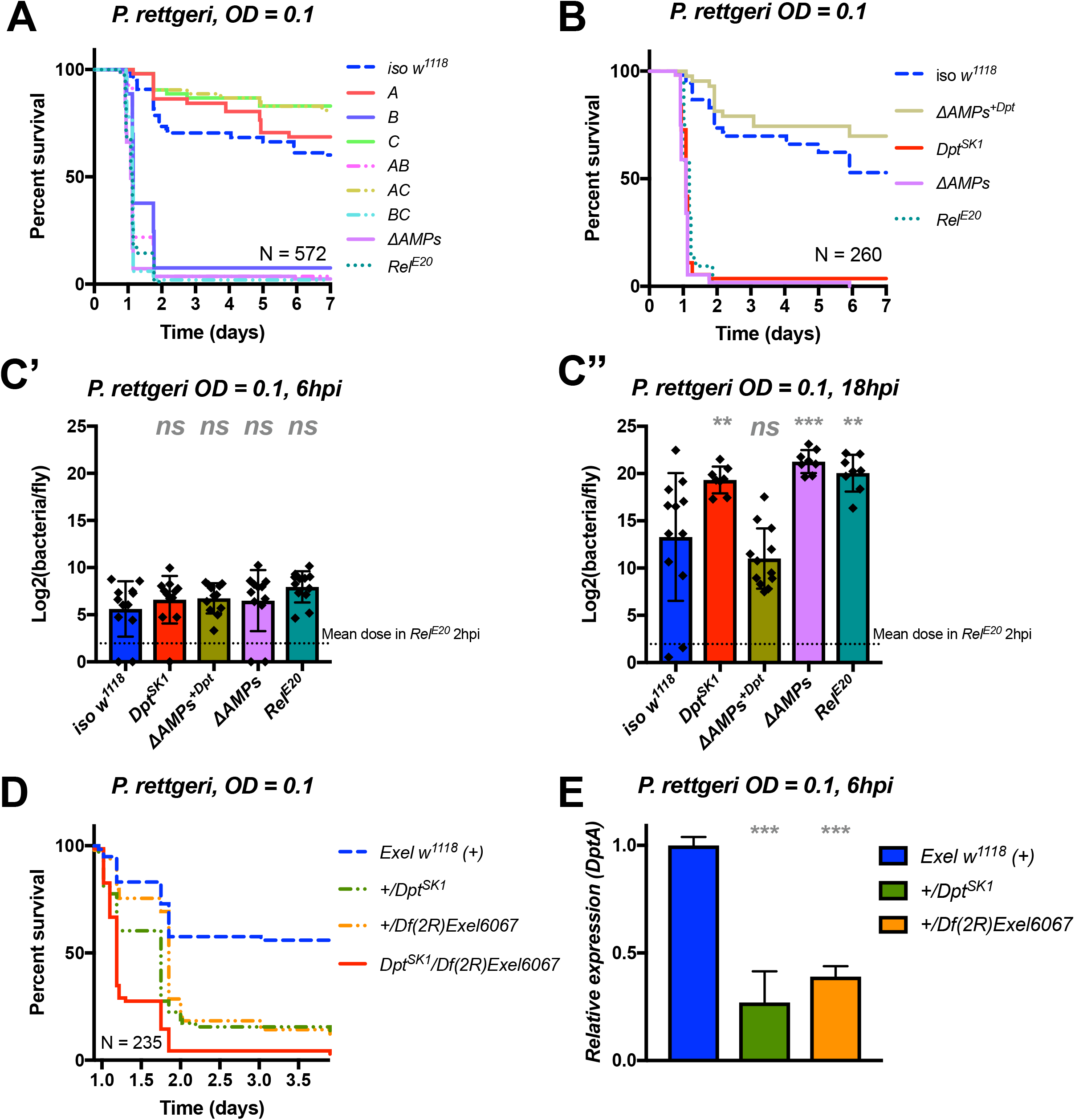
Identification of AMPs involved in the susceptibility of *ΔAMPs* flies to *P. rettgeri.* A) Survival of mutants for groups of AMPs reveals that only loss of Imd-responsive Group B peptides (Drosocin, Attacins, and Diptericins) recapitulates the susceptibility of *ΔAMPs* flies. B) Further dissection of the mutations affected in Group B reveals that only the loss of Diptericins (*Dpt*^SK1^) leads to susceptibility similar to *ΔAMPs* flies. Remarkably, flies lacking all other AMPs *(ΔAMPs+^Dpt^)* resist as wild-type. C) Bacterial loads of individual flies are similar at 6hpi (C’), but by 18hpi (C”), *Dpt* mutants and *Rel^E20^* flies have all failed to control *P. rettgeri* growth. D) Heterozygote flies for *Dpt^SK1^* and a deficiency including the *Diptericins* and flanking genes *(Df(2R)Exel6067)* recapitulates the susceptibility of *Diptericin* mutants. Intriguingly, heterozygotes with one functional copy of the Diptericins *(+/Dpt^SK1^* or *+/ Df(2R)Exel6067)* are nonetheless highly susceptible to infection. E) *Diptericin A* transcriptional output is strongly reduced in heterozygotes 6hpi compared to wild-type flies. One-way ANOVA: not significant = *ns,* p < .05 = *, p < .01 = **, and p < .001 = *** relative to *iso w^1118^*.

Collectively, our study shows that *Diptericins* are critical to resist *P. rettgeri,* while they play an important but less essential role in defence against *P. burhodogranariea* infection. We were curious whether *Diptericin’s* major contribution to defence observed with *P. rettgeri* could be generalized to other members of the genus *Providencia.* An exclusive role for *Diptericins* was also found for the more virulent *P. stuartii* (Fig. 5 supplement C), but not for other *Providencia* species tested (P. *burhodogranariea, P. alcalifaciens, P. sneebia, P. vermicola)* (data not shown).

### *Drosocin* is critical to resist infection with *E. cloacae*

In the course of our exploration of AMP-pathogen interactions, we identified another highly specific interaction between *E. cloacae* and Drosocin. Use of compound mutants revealed that alone, Group B flies were already susceptible to *E. cloacae.* Meanwhile, Group AB flies reached *ΔAMPs* levels of susceptibility, while Group A and Group C flies resisted as wild-type (Fig. 6A). The high susceptibility of Group AB flies results from a synergistic interaction amongst Group A and Group B peptides in defence against *E. cloacae (A*B,* HR = +2.55, p = .003).

**Figure 6:**
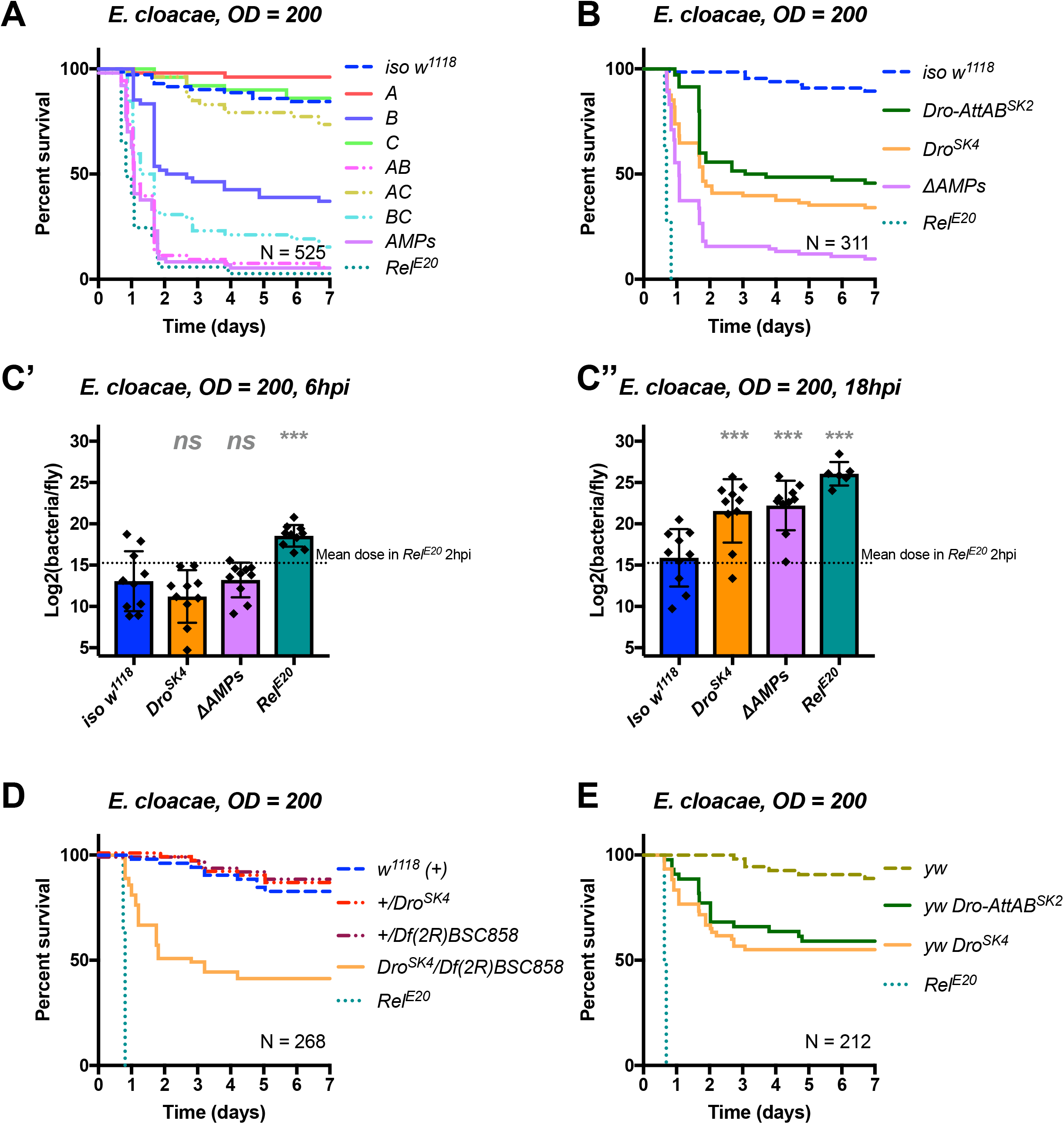
Identification of AMPs involved in the susceptibility of *ΔAMPs* flies to *E. cloacae.* A) Survival of mutants for groups of AMPs reveals that loss of Imd-responsive Group B peptides (Drosocin, Attacins, and Diptericins) results in a strong susceptibility to infection (p < .001), while loss of Group A or C peptides alone resists as wild-type (p > 0.1 each). Group AB flies were as susceptible as *ΔAMPs* flies, and we observed a synergistic interaction between Group A and B mutations (A*B: HR = +2.55, p = .003). B) Further dissection of the mutations in Group B revealed that loss of *Drosocin* alone *(Dro^SK4^),* or a deficiency lacking both *Drosocin* and *Attacins AttA* and *AttB (Dro-AttAB^SK2^)* recapitulates the susceptibility of Group B flies. C) By 18hpi, bacterial loads in individual *Drosocin* mutants or *Rel^E20^* flies are significantly higher than wild-type. D) Heterozygote flies for *Dro^SK4^* and *Df(2R)BSC858* (a deficiency removing *Drosocin, Attacins AttA* and *AttB,* and other genes) are strongly susceptible to *E. cloacae* infection. E) *Drosocin* mutants in an alternate genetic background *(yw*) are susceptible to *E. cloacae.* One-way ANOVA: not significant = *ns,* and p < .001 = *** relative to *iso w^1118^*.

We chose to further explore the AMPs deleted in Group B flies, as alone this genotype already displayed a strong susceptibility. Use of individual mutant lines revealed that mutants for *Drosocin* alone *(Dro^SK4^)* or the *Drosocin-Attacin A/B* deficiency *(Dro-AttAB^SK2^),* but not *AttC, AttD, nor Dpt^SK1^* (not shown), recapitulate the susceptibility observed in Group B flies (Fig. 6B). At 18hpi, both *Dro^SK4^* and *ΔAMPs* flies had significantly higher bacterial loads compared to wild-type flies, while *Rel^E20^* mutants were already moribund with much higher bacterial loads (Fig. 6C). Indeed, the deletion of *Drosocin* alone drastically alters the fly’s ability to control the otherwise avirulent *E. cloacae* with inoculations at OD=200 (~39,000 bacteria, Fig. 6A-C) or even OD=10 (~7,000 bacteria, Fig. 6 supplement A).

We confirmed the high susceptibility of *Drosocin* mutant flies to *E. cloacae* in various contexts: transheterozygote flies carrying *Dro^SK4^* over a *Drosocin* deficiency *(Df(2R)BSC858)* that also lacks flanking genes including *AttA* and *AttB* ((Fig. 6D), the *Dro^SK4^* mutations in an alternate genetic background *(yw*, Fig. 6E), and, *Drosocin RNAi* (Fig. 6 supplement B,C). Thus, we recovered two highly specific AMP-pathogen interactions: Diptericins are essential to combat *P. rettgeri* infection, while *Drosocin* is paramount to surviving *E. cloacae* infection.

## Discussion

### A combinatory approach to study AMPs

Despite the recent emphasis on innate immunity, little is known on how immune effectors contribute individually or collectively to host defence, exemplified by the lack of in depth *in vivo* functional characterization of *Drosophila* AMPs. Taking advantage of new gene editing approaches, we developed a systematic mutation approach to study the function of *Drosophila* AMPs. With eight distinct mutations, we were able to generate a fly line lacking the 14 AMPs that are inducible during the systemic immune response. A striking first finding is that *ΔAMPs* flies were perfectly healthy and have an otherwise wild-type immune response. This indicates that in contrast to mammals^15^, *Drosophila* AMPs are not likely to function as signaling molecules. Most flies lacking a single AMP family exhibited a higher susceptibility to certain pathogens consistent with their *in vitro* activity. We found activity of Diptericins against *P. rettgeri,* Drosocin against *E. cloacae,* Drosomycin and Metchnikowin against *C. albicans,* and Defensin and Cecropin against *P. burhodogranariea* (Fig. 4 supplement A). In most cases, the susceptibility of single mutants was slight, and the contribution of individual AMPs could be revealed only when combined to other AMP mutations as illustrated by the susceptibility of *Drosocin, Attacin,* and *Diptericin* combined mutants to *P. burhodogranariea.* Thus, the use of compound rather than single mutations provides a better strategy to decipher the contribution of AMPs to host defence.

### AMPs and Bomanins are essential contributors to Toll and Imd pathway mediated host defence

The Toll and Imd pathways provide a paradigm of innate immunity, illustrating how two distinct pathways link pathogen recognition to distinct but overlapping sets of downstream immune effectors^1,63^. However, a method of deciphering the contributions of the different downstream effectors to the specificity of these pathways remained out of reach, as mutations in these immune effectors were lacking. Our study shows that AMPs contribute greatly to resistance to Gram-negative bacteria. Consistent with this, *ΔAMPs* flies are almost as susceptible as Imd-deficient mutants to most Gram-negative bacteria. In contrast, flies lacking AMPs were only slightly more susceptible to Gram-positive bacteria and fungal infections compared to wild-type flies, and this susceptibility rarely approached the susceptibility of *Bomanin* mutants. This may be due to the cell walls of Gram-negative bacteria being thinner and more fluid than the rigid cell walls of Gram-positive bacteria^64^, consequently making Gram-negative bacteria more prone to the action of pore-forming cationic peptides. It would be interesting to know if the specificity of AMPs to primarily combatting Gram-negative bacteria is also true in other species.

Based on our study and Clemmons et al.^45^, we can now explain the susceptibility of Toll and Imd mutants at the level of the effectors, as we show that mutations affecting Imd-pathway responsive antibacterial peptide genes are highly susceptible to Gram-negative bacteria while the Toll-responsive targets Drosomycin, Metchnikowin, and especially the Bomanins, confer resistance to fungi and Gram-positive bacteria. Thus, the susceptibility of these two pathways to different sets of microbes not only reflects specificity at the level of recognition, but can now also be translated to the activities of downstream effectors. It remains to be seen how Bomanins contribute to the microbicidal activity of immune-induced hemolymph, as attempts to synthesize Bomanins have not revealed direct antimicrobial activity^46^. It should also be noted that many putative effectors downstream of Toll and Imd remain uncharacterized, and so could also contribute to host defence beyond AMPs and Bomanins.

### AMPs act additively and synergistically to suppress bacterial growth in vivo

In the last few years, numerous *in vitro* studies have focused on the potential for synergistic interactions of AMPs in microbial killing^7,54,56,65–70^. Our collection of AMP mutant fly lines placed us in an ideal position to investigate AMP interactions in an *in vivo* setting. While Toll-responsive AMPs (Group C: Metchnikowin, Drosomycin) additively contributed to defence against the yeast *C. albicans,* we found that certain combinations of AMPs have synergistic contributions to defence against *P. burhodogranariea.* Synergistic loss of resistance may arise in two general fashions: first, co-operation of AMPs using similar mechanisms of action may breach a threshold microbicidal activity whereupon pathogens are no longer able to resist. This may be the case for our observations of synergy amongst Diptericins and Attacins against *P. burhodogranariea,* as only co-occurring loss of both these related glycine-rich peptide families^36^ led to complete loss of resistance. Alternatively, synergy may arise due to complementary mechanisms of action, whereupon one AMP potentiates the other AMP’s ability to act. For instance, the action of the bumblebee AMP Abaecin, which binds to the molecular chaperone DnaK to inhibit bacterial DNA replication, is potentiated by the presence of the pore-forming peptide Hymenoptaecin^71^. *Drosophila* Drosocin is highly similar to Abaecin, including O-glycosylation of a critical threonine residue^2,72^, and thus likely acts in a similar fashion. Furthermore, *Drosophila* Attacin C is maturated into both a glycine-rich peptide and a Drosocin-like peptide called MPAC^73^. As such, co-occuring loss of Drosocin, MPAC, and other possible MPAC-like peptides encoded by the Attacin/Diptericin superfamily may be responsible for the synergistic loss of resistance in *Drosocin, Attacin, Diptericin* combined mutants.

### AMPs can act with great specificity against certain pathogens

It is commonly thought that the innate immune response lacks the specificity of the adaptive immune system, which mounts directed defences against specific pathogens. Accordingly for innate immunity, the diversity of immune-inducible AMPs can be justified by the need for generalist and/or co-operative mechanisms of microbial killing. However, an alternate explanation may be that innate immunity expresses diverse AMPs in an attempt to hit the pathogen with a “silver bullet:” an AMP specifically attuned to defend against that pathogen. Here, we provide a demonstration in an *in vivo* setting that such a strategy may actually be employed by the innate immune system. Remarkably we recovered not just one, but two examples of exquisite specificity in our laborious but relatively limited assays.

*Diptericin* has previously been highlighted for its important role in defence against *P. rettgeri^62^,* but it was previously unknown whether other AMPs may confer defence in this infection model. Astoundingly, flies mutant for all other inducible AMPs resisted *P. rettgeri* infection as wild-type, while only *Diptericin* mutants succumbed to infection. This means that Diptericins do not co-operate with other AMPs in defence against *P. rettgeri*, and are solely responsible for defence in this specific host-pathogen interaction. Moreover, *+/Dpt^SK1^* heterozygote flies were nonetheless extremely susceptible to infection, demonstrating that a full transcriptional output over the course of infection is required to effectively prevent pathogen growth. A previous study has shown that ~7hpi appears to be the critical time point at which *P. rettgeri* either grows unimpeded or the infection is controlled^24^. This time point correlates with the time at which the *Diptericin* transcriptional output is in full-force^41^. Thus, a lag in the transcriptional response in *Dpt^SK1^/+* flies likely prevents the host from reaching a competent Diptericin concentration, indicating that *Diptericin* expression level is a key factor in successful host defence.

We also show that *Drosocin* is specifically required for defence against *E. cloacae.* This striking finding validates previous biochemical analyses showing Drosocin *in vitro* activity against several Enterobacteriaceae, including *E. cloacae^37^.* As *ΔAMPs* flies are more susceptible than *Drosocin* single mutants, other AMPs also contribute to Drosocin-mediated control of *E. cloacae.* As highlighted above, Drosocin is similar to other Proline-rich AMPs (*e.g.* Abaecin, Pyrrhocoricin) that have been shown to target bacterial DnaK^6,7^. Alone, these peptides still penetrate bacteria cell walls through their uptake by bacterial permeases^71,74^. Thus, while Drosocin would benefit from the presence of pore-forming toxins to enter bacterial cells^71^, the veritable “stake to the heart” is likely the plunging of Drosocin itself into vital bacterial machinery.

### On the role of AMPs in host defence

It has often been questioned why flies should need so many AMPs^1,4,75^. A common idea, supported by *in vitro* experiments^7,65,70^, is that AMPs work as cocktails, wherein multiple effectors are needed to kill invading pathogens. However, we find support for an alternative hypothesis that suggests AMP diversity may be due to highly specific interactions between AMPs and subsets of pathogens that they target. Burgeoning support for this idea also comes from recent evolutionary studies that show *Drosophila* and vertebrate AMPs experience positive selection^62,72,75–81^, a hallmark of host-pathogen evolutionary conflict. Our functional demonstrations of AMP-pathogen specificity, using naturally relevant pathogens^60,82^, suggest that such specificity is fairly common, and that certain AMPs can act as the arbiters of life or death upon infection by certain pathogens. This stands in contrast to the classical view that the AMP response contains such redundancy that single peptides should have little effect on organism-level immunity^4,61,75,83^. Nevertheless, it seems these immune effectors play non-redundant roles in defence.

By providing a long-awaited *in vivo* functional validation for the role of AMPs in host defence, we also pave the way for a better understanding of the functions of immune effectors. Our approach of using multiple compound mutants, now possible with the development of new genome editing approaches, was especially effective to decipher the logic of immune effectors. Understanding the role of AMPs in innate immunity holds great promise for the development of novel antibiotics^18,84,85^, insight into autoimmune diseases^86–89^, and given their potential for remarkably specific interactions, perhaps in predicting key parameters that predispose individuals or populations to certain kinds of infections^61,75,76^. Finally, our set of isogenized *AMP* mutant lines provides long-awaited tools to decipher the role of AMPs not only in immunity, but also in the various roles that AMPs may play in aging, neurodegeneration, anti-tumour activity, regulation of the microbiota and more, where disparate evidence has pointed to their involvement.

**Figure 1 supplement:**
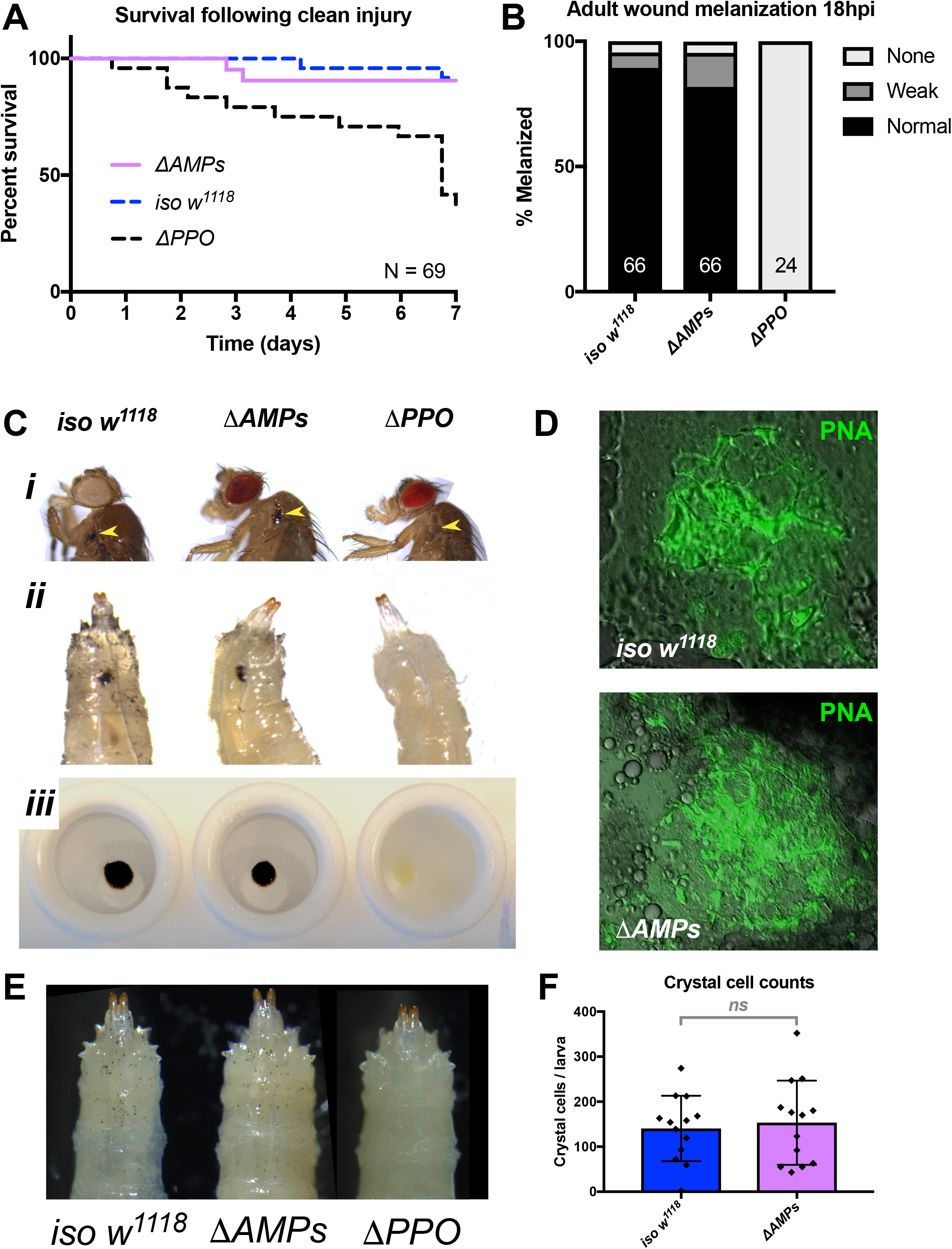
*ΔAMPs* flies have otherwise wild-type immune reactions. A) *ΔAMPs* flies survive clean injury like wild-type flies, while *ΔPPO* mutants deficient for melanization have reduced survival over time. B) *ΔAMPs* flies melanize the cuticle similar to wild-type flies following pricking (χ^2^ = 2.14, p = .34). Melanization categories (None, Weak, Normal) were as described in Dudzic et al.^90^. Sample sizes (n) are included in each bar. C) Melanization in *iso w^1118^, ΔAMPs,* and *ΔPPO* flies of the cuticle in adults (*i*, yellow arrowheads), larvae *(ii,* melanized wounds), and larval hemolymph (iii). D) To investigate clotting ability, we used the hanging drop assay^50^ with *ΔAMPs* larval hemolymph and visualized clot fibers with PNA staining (green). Both *iso w^1118^* and *ΔAMPs* hemolymph produced visible clot fibres measured after 20 minutes. Hemocyte populations are normal in *ΔAMPs* flies, including crystal cell distribution (E) and number (F).

**Figure 2 supplement:**
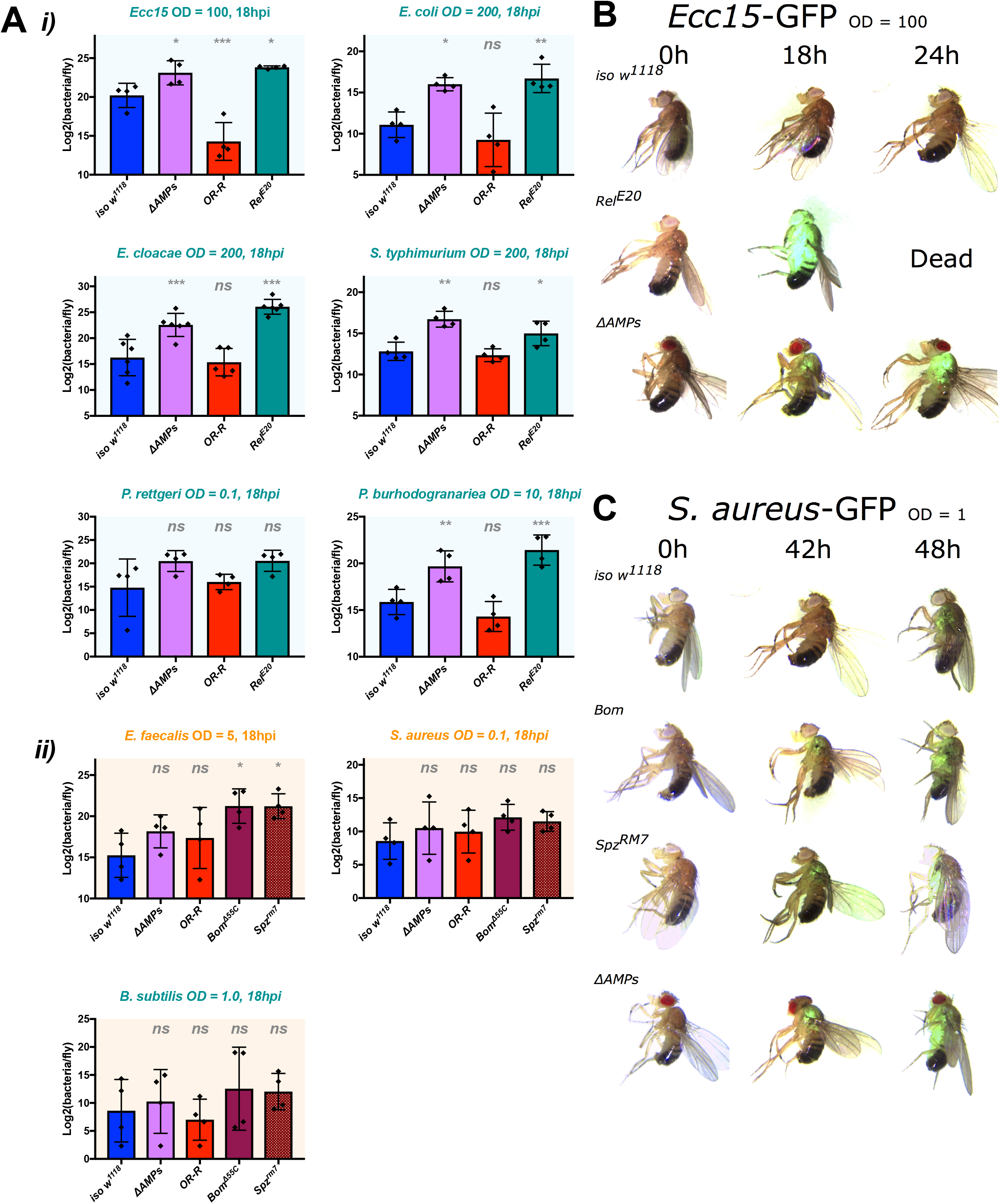
*ΔAMPs* flies fail to suppress Gram-negative bacterial growth. Colony counts were performed on pooled samples (5 flies) for bacteria amenable to LB agar, a medium that avoids overnight growth of the host microbiota. A) For Gram-negative bacterial infections, *ΔAMPs* flies have significantly higher bacterial loads than *iso w^1118^* at 18 hours post-infection (hpi) (i). This is not true for any of the Gram-positive bacteria tested (ii), while *spz^rm7^* mutants carried higher bacterial loads, significantly so in *E. faecalis* infections. Gram-negative (B) and Gram-positive (C) infections with GFP-labelled bacteria spread from the wound site systemically in all genotypes tested. Thus *ΔAMPs* fly mortality is likely not due to tissue-specific colonization by invading bacteria, but rather a failure to suppress bacterial growth first locally, and then systemically. One-way ANOVA: not significant = *ns*, p < .05 = *, p < .01 = **, and p < .001 = *** relative to *iso w^1118^*.

**Figure 4 supplement:**
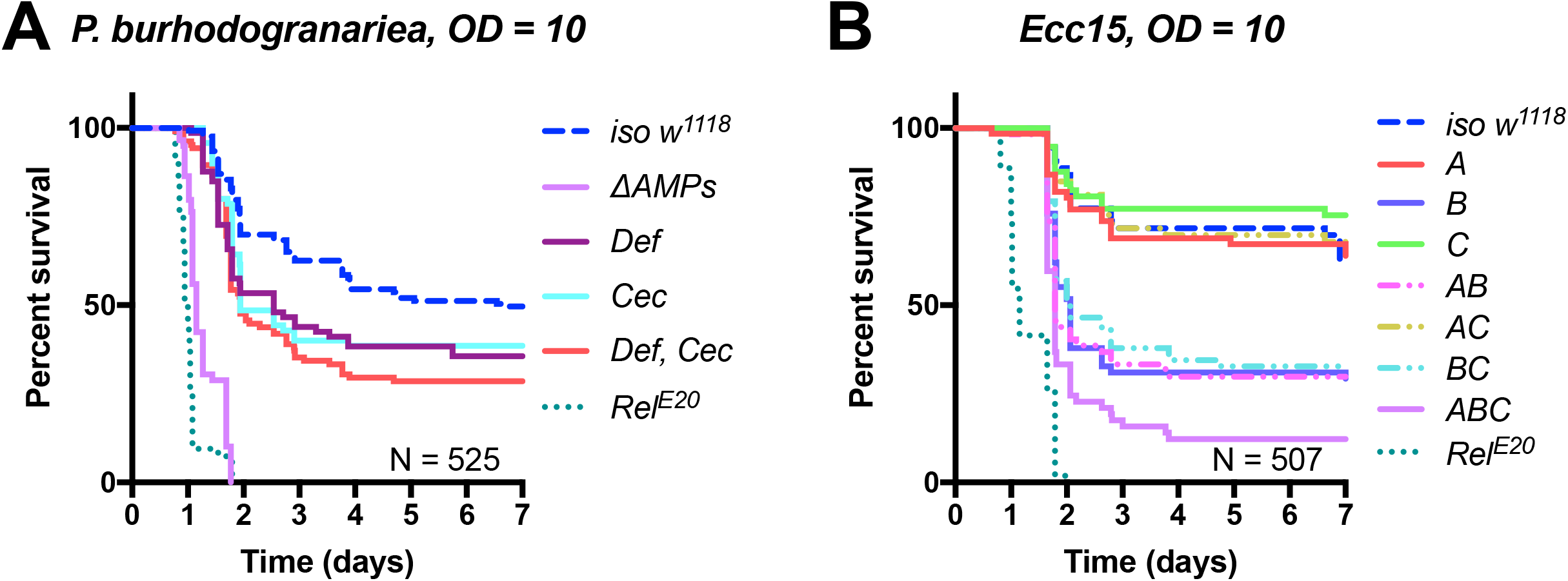
A) Dissection of the susceptibility of Group A flies lacking *Defensin* and *Cecropins* reveals that combined mutants have an additive loss of resistance *(Def*Cec,* HR = +0.36, p = .342). B) Upon infection with the Gram-negative *Ecc15,* Group B peptides (Drosocin, Attacins and Diptericins) explain the bulk of mortality, but additional loss of other peptides in *ΔAMPs* flies leads to increased mortality (Log-Rank p = .013).

**Figure 5 supplement:**
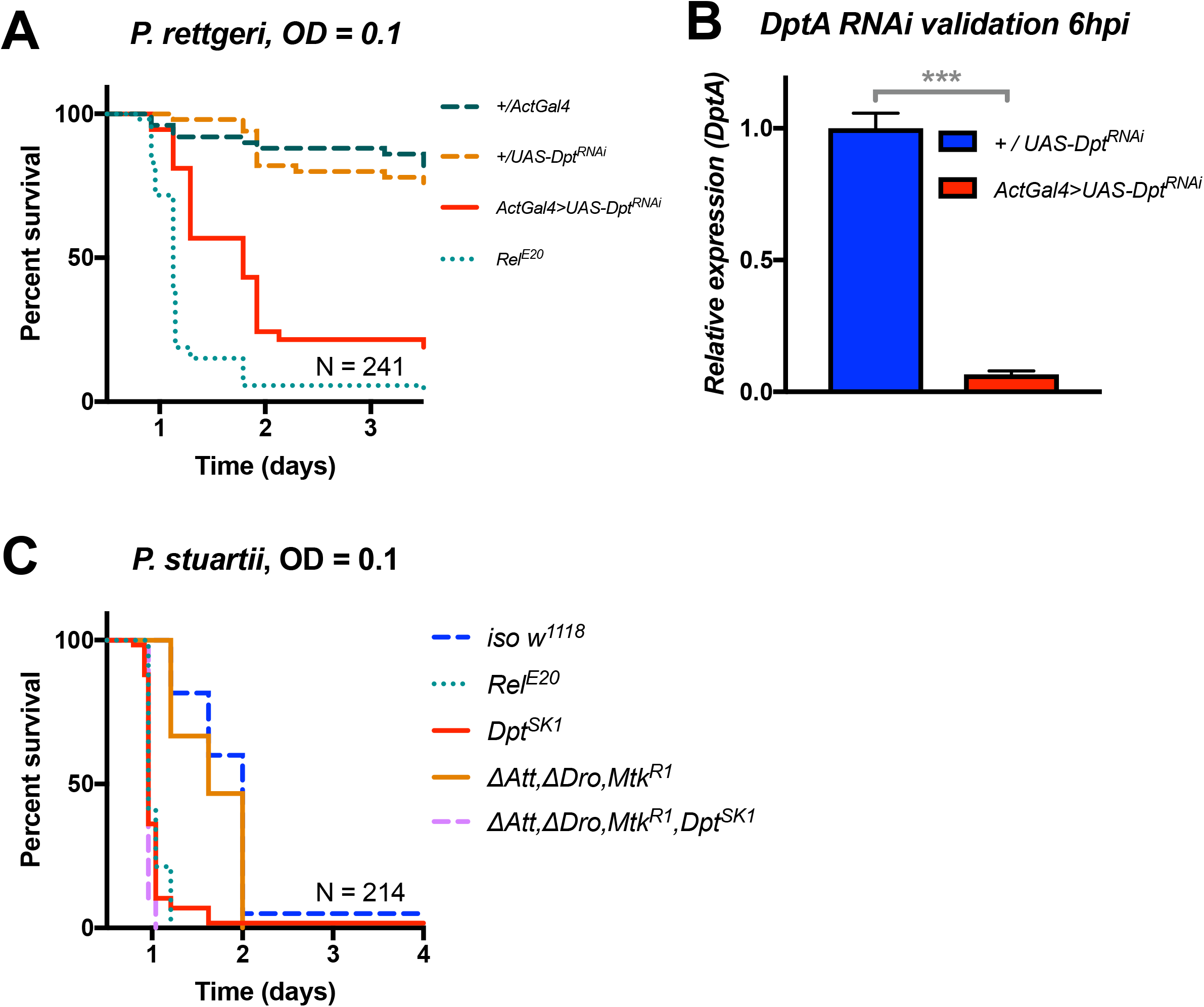
A) Silencing of *Diptericin by RNAi* leads to higher susceptibility to *P. rettgeri* infection (p < .001). B) Validation of the *Diptericin* RNAi construct 6hpi. C) Mutants lacking multiple peptides (Attacins, Drosocin, and Metchnikowin) succumb to *P. stuartii* infection as wild-type (‘ΔAtt, *ΔDro, Mtk^R1f^*), while Diptericin mutation alone *(Dpt^SK1^)* or combined (‘*ΔAtt, ΔDro, Mtk^R1^, ΔDpt’*) leads to a susceptibility similar to *Rel^E20^* mutants. This pattern of survival was similar to the pattern observed with *P. rettgeri.* One-way ANOVA: p < .001 = ***.

**Figure 6 supplement:**
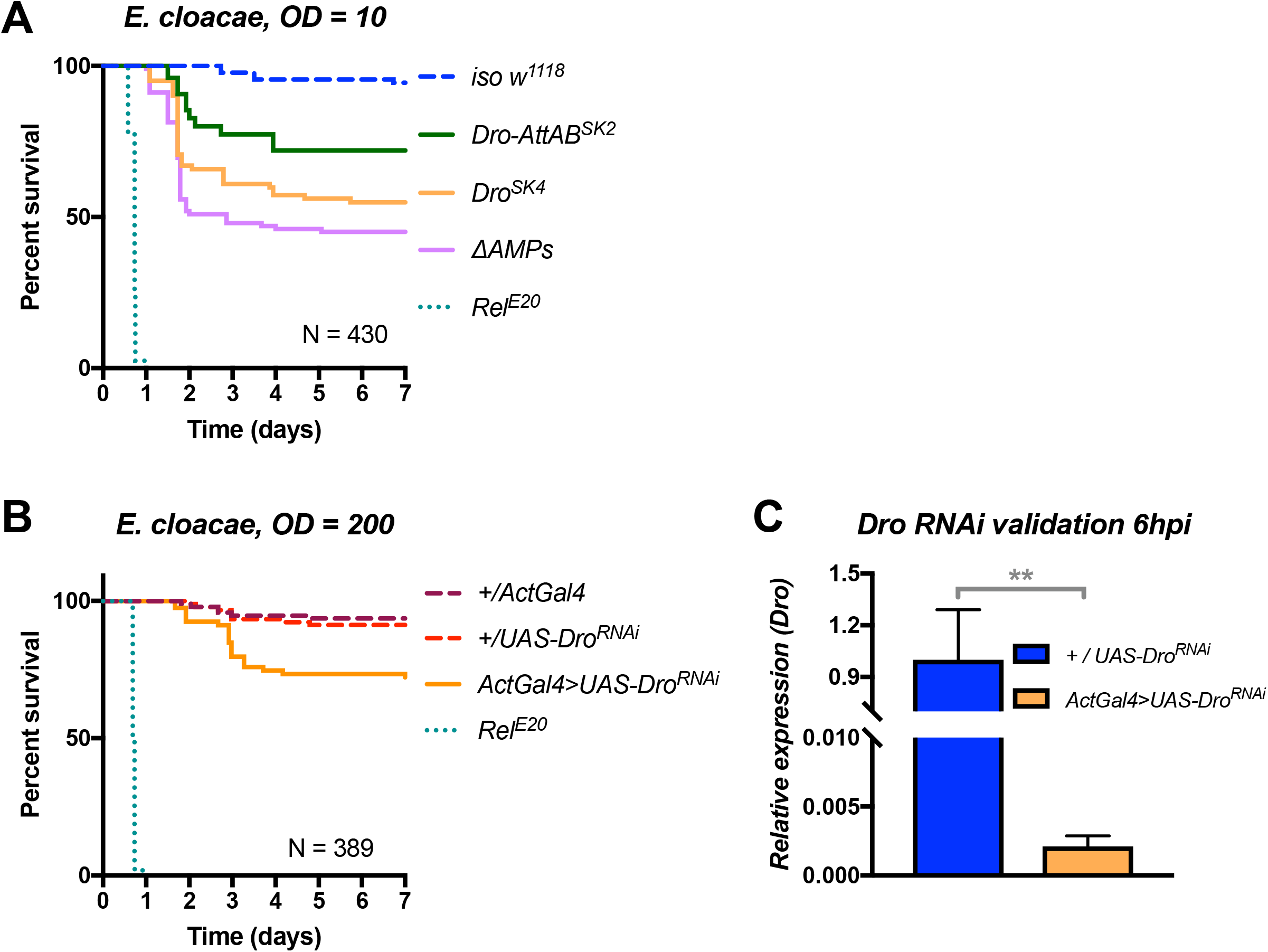
A) *Drosocin* mutant susceptibility remains even at a lower dose (OD=10, ~7000 bacteria/fly), while *Rel^E20^* flies succumb rapidly regardless of initial dose. B) Silencing of *Drosocin by RNAi* leads to significant mortality from *E. cloacae* infection (p < .001). C) Validation of the *Drosocin* RNAi construct 6hpi.

**Figure S1:**
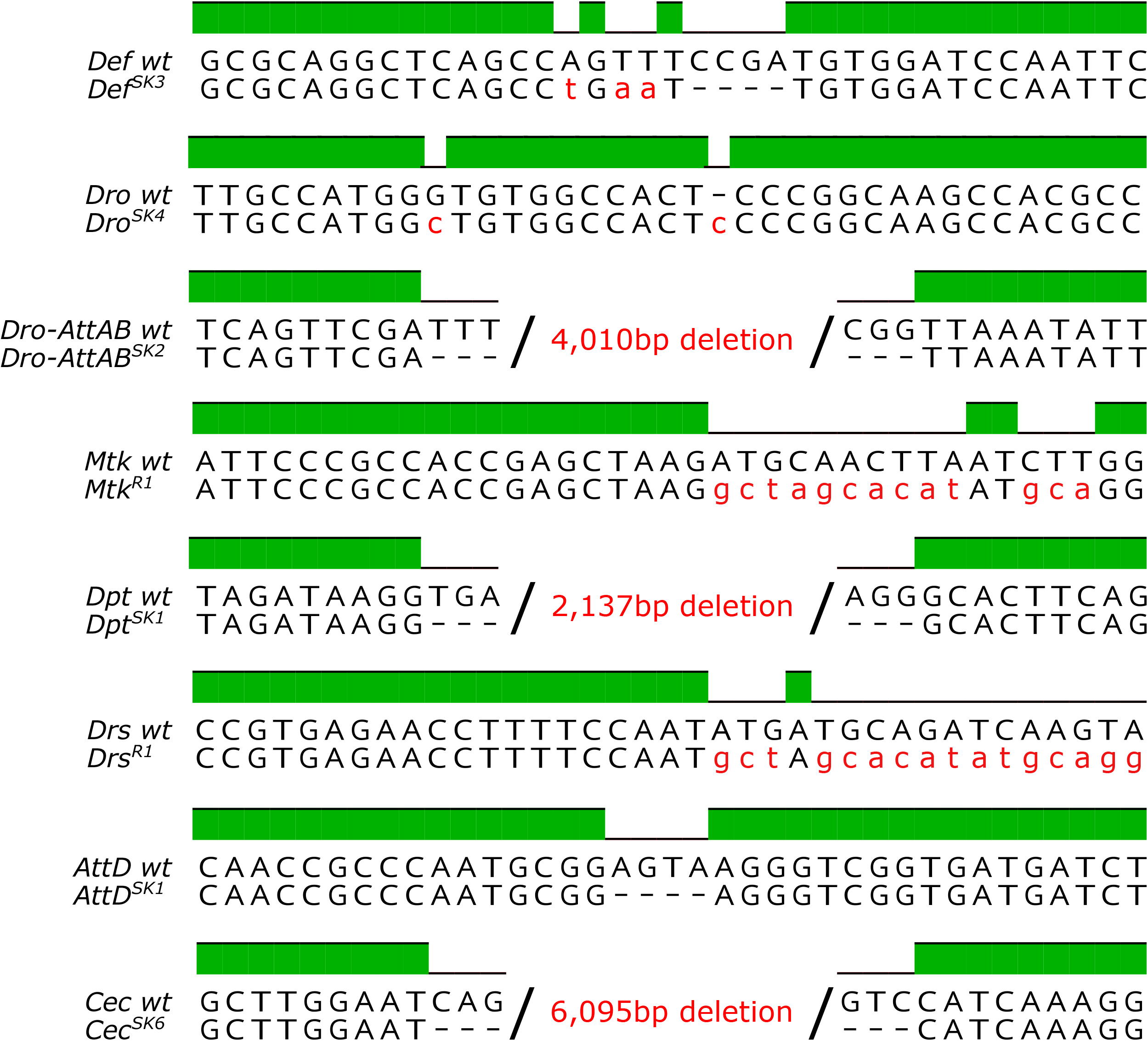
Genetic description of mutations generated in this study. *Mtk^R1^* and *Drs^R1^* mutations entirely replaced the CDS with an insert from the piHR vector. Non-synonymous nucleotides in mutants are given in red.

**Table S1:**
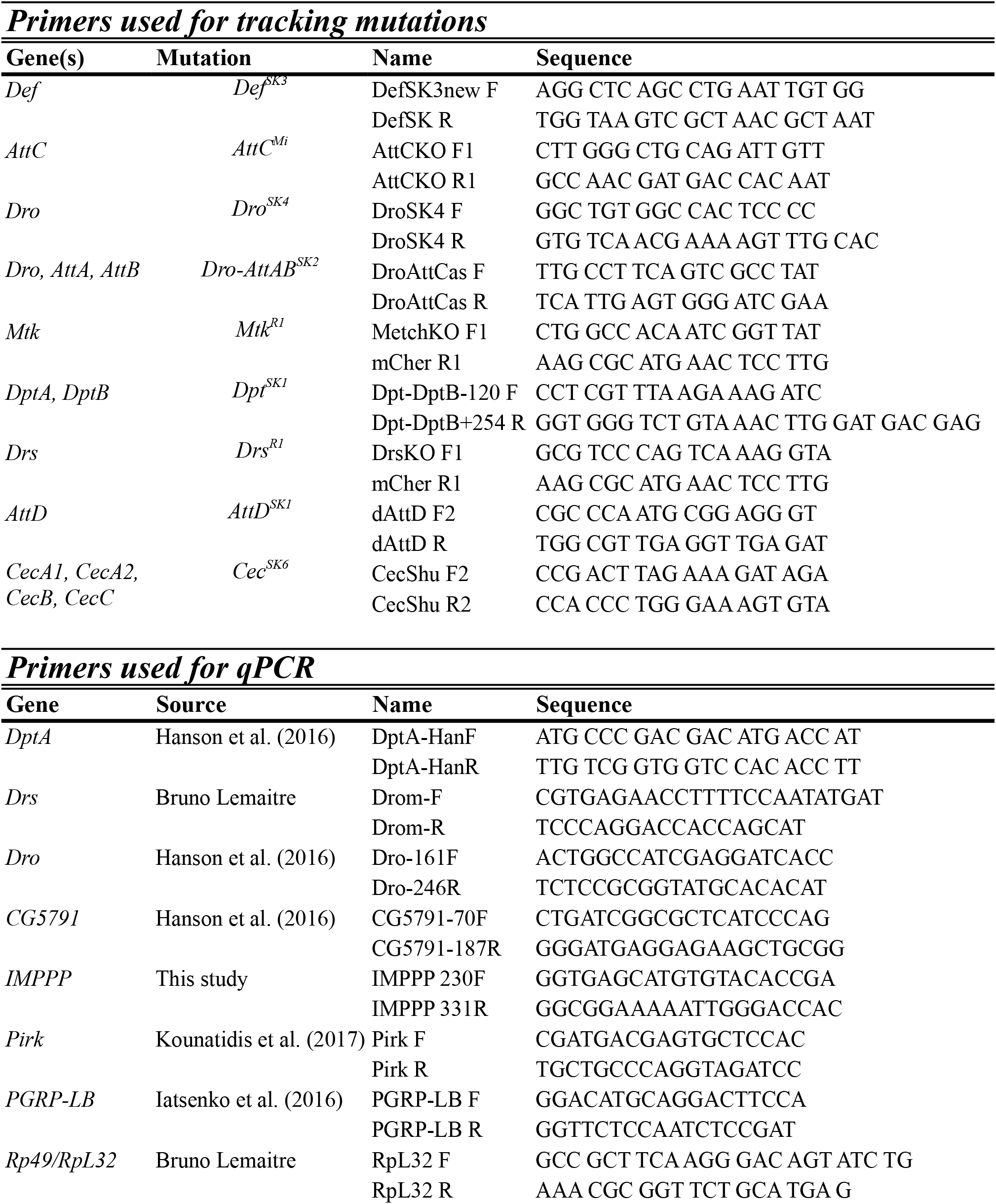
Primers used in this study to track AMP mutations or measure gene expression.

## Materials and Methods

### Drosophila genetics and mutant generation

The DrosDel^48^ isogenic *w^1118^ (iso w^1118^)* wild type was used as a genetic background for mutant isogenization. Alternate wild-types used throughout include Oregon R (OR-R), *w^1118^* from the Vienna Drosophila Resource Centre, and the Canton-S isogenic line Exelexis *w^1118^*, which was kindly provided by Brian McCabe. *Bom^Δ55C^* mutants were generously provided by Steven Wasserman, and *Bom^Δ55C^* was isogenized into the *iso w^1118^*background. *Rel^E20^* and *spz^rm7^ iso w^1118^* flies were provided by Luis Teixeira^51,91^. Prophenoloxidase mutants *(ΔPPO)* are described in Dudzic et al.^92^. P-element mediated homologous recombination according to Baena-Lopez et al.^93^ was used to generate mutants for *Mtk (Mtk^R1^)* and *Drs* (Drs^R1^).

Plasmids were provided by Mickael Poidevin. *Attacin C*mutants *(AttC^Mi^,* #25598), the *Diptericin* deficiency *(Df(2R)Exel6067,* #7549), the *Drosocin* deficiency *(Df(2R)BSC858,* #27928), *UAS-Diptericin RNAi (Dpt^RNAi^,* #53923), *UAS-Drosocin RNAi (Dro^RNAi^,* #67223), and *Actin5C-Gal4 (ActGal4,* #4414) were ordered from the Bloomington stock centre (stock #s included). CRISPR mutations were performed by Shu Kondo according to Kondo and Ueda^94^, and full descriptions are given in Figure S1. In brief, flies deficient for *Drosocin, Attacin A,* and *Attacin B (Dro-AttAB^SK2^), Diptericin A* and *Diptericin B (Dpt^SK1^),* and *Cecropins CecA1, CecA2, CecB, CecC (Cec^SK6^)* were all produced by gene region deletion specific to those AMPs without affecting other genes. Single mutants for *Defensin (Def^SK3^), Drosocin (Dro^SK4^),* and *Attacin D (AttD^SK1^)* are small indels resulting in the production of short (80-107 residues) nonsense peptides. Mutations were isogenized for a minimum of seven generations into the *iso w^1118^* background prior to subsequent recombination.

### Microbial culture conditions

Bacteria were grown overnight on a shaking plate at 200rpm in their respective growth media and temperature conditions, and then pelleted by centrifugation at 4°C. These bacterial pellets were diluted to the desired optical density at 600nm (OD) as indicated. The following bacteria were grown at 37°C in LB media: *Escherichia coli strain 1106, Salmonella typhimurium, Enterobacter cloacae ß12, Providencia rettgeri strain Dmel, Providencia burhodogranariea strain B, Providencia stuartii strain DSM 4539, Providencia sneebia strain Dmel, Providencia alcalifaciens strain Dmel, Providencia vermicola strain DSM 17385, Bacillus subtilis,* and *Staphylococcus aureus. Erwinia carotovora carotovora (Ecc15)* and *Micrococcus luteus* were grown overnight in LB at 29°C. *Enterococcus faecalis* and *Listeria innocua* were cultured in BHI medium at 37°C. *Candida albicans* strain ATCC 2001 was cultured in YPG medium at 37°C. *Aspergillus fumigatus* was grown at room temperature on Malt Agar, and spores were collected in sterile PBS rinses, pelleted by centrifugation, and then resuspended to the desired OD in PBS. The entomopathogenic fungi *Beauveria bassiana* and *Metarhizium anisopliae* were grown on Malt Agar at room temperature until sporulation.

### Systemic infections and survival

Systemic infections were performed by pricking 3-5 day old adult males in the thorax with a 100 μm thick insect pin dipped into a concentrated pellet of bacteria or fungal spores. Infected flies were subsequently maintained at 25°C for experiments. For infections with *B. bassiana* and *M. anisopliae,* flies were anaesthetized and then shaken on a sporulating plate of fungi for 30s. At least two replicate survival experiments were performed for each infection, with 20-35 flies per vial on standard fly medium without yeast. Survivals were scored twice daily, with additional scoring at sensitive time points. Comparisons of *iso w^1118^* wild-type to *ΔAMPs* mutants were made using a Cox-proportional hazard (CoxPH) model, where independent experiments were included as covariates, and covariates were removed if not significant (p > .05). Direct comparisons were performed using Log-Rank tests in Prism 7 software. The effect size and direction is included as the CoxPH hazard ratio (HR) where relevant, with a positive effect indicating increased susceptibility. CoxPH models were used to test for synergistic contributions of AMPs to survival in R 3.4.4. Total sample size (N) is given for each experiment as indicated.

### Quantification of microbial load

The native *Drosophila* microbiota does not readily grow overnight on LB, allowing for a simple assay to estimate bacterial load. Flies were infected with bacteria at the indicated OD as described, and allowed to recover. At the indicated time postinfection, flies were anaesthetized using CO_2_ and surface sterilized by washing them in 70% ethanol. Ethanol was removed, and then flies were homogenized using a Precellys™ bead beater at 6500rpm for 30 seconds in LB broth, with 300ul for individual samples, or 500uL for pools of 5-7 flies. These homogenates were serially diluted and 150uL was plated on LB agar. Bacterial plates were incubated overnight, and colony-forming units (CFUs) were counted manually. Statistical analyses were performed using One-way ANOVA with Sidak’s correction. P-values are reported as < 0.05 = *, < 0.01 = **, and < 0.001 = ***. For *C. albicans,* BiGGY agar was used instead to select for *Candida* colonies from fly homogenates.

### Gene expression by qPCR

Flies were infected by pricking flies with a needle dipped in a pellet of either *E. coli* or *M. luteus* (OD600 = 200), and frozen at -20°C 6h and 24h post-infection respectively. Total RNA was then extracted from pooled samples of five flies each using TRIzol reagent, and re-suspended in MilliQ dH_2_O. Reverse transcription was performed using 0.5 micrograms total RNA in 10 μl reactions using PrimeScript RT (TAKARA) with random hexamer and oligo dT primers. Quantitative PCR was performed on a LightCycler 480 (Roche) in 96-well plates using Applied Biosystems™ SYBR™ Select Master Mix. Values represent the mean from three replicate experiments. Error bars represent one standard deviation from the mean. Primers used in this study can be found in Table S1. Statistical analyses were performed using one-way ANOVA with Tukey post-hoc comparisons. P-values are reported as not significant = ns, < 0.05 = *, < 0.01 = **, and < 0.001 = ***. qPCR primers and sources^11,72,95^ are included in Table S1.

### MALDI-TOF peptide analysis

Two methods were used to collect hemolymph from adult flies: in the first method, pools of five adult females were pricked twice in the thorax and once in the abdomen. Wounded flies were then spun down with 15μL of 0.1% trifluoroacetic acid (TFA) at 21000 RCF at 4°C in a mini-column fitted with a 10μm pore to prevent contamination by circulating hemocytes. These samples were frozen at -20°C until analysis, and three biological replicates were performed with 4 technical replicates. In the second method, approximately 20nL of fresh hemolymph was extracted from individual adult males using a Nanoject, and immediately added to 1μL of 1% TFA, and the matrix was added after drying. Peptide expression was visualized as described in Üttenweiller-Joseph et al.^49^. Both methods produced similar results, and representative expression profiles are given.

### Melanization and hemocyte characterization, image acquisition

Melanization assays^90^ and peanut agglutinin (PNA) clot staining^50^ was performed as previously described. In brief, flies or L3 larvae were pricked, and the level of melanization was assessed at the wound site. We used FACS sorting to count circulating hemocytes. For sessile crystal cell visualization, L3 larvae were cooked in dH_2_O at 70°C for 20 minutes, and crystal cells were visualized on a Leica DFC300FX camera using Leica Application Suite and counted manually.

## Author contributions

MAH, AD and BL designed the study. MAH and AD performed DrosDel isogenization and recombination. MP and SK supplied critical reagents. MAH performed the experiments, and CC provided experimental support. MAH and BL analyzed the data and wrote the manuscript. All authors read and approved the final manuscript.

## Acknowledgements

We would like to thank Marc Moniatte for assistance with MALDI-TOF analysis, Claudia Melcarne for assistance with hemocyte characterization, and Igor Iatsenko for help in preparation of critical reagents. Brian Lazzaro generously provided *Providencia* species used in this study. We would like to thank Hannah Westlake for useful comments on the manuscript. MAH would like to extend special thanks to Jan Dudzic for many illuminating discussions had over coffee.

